# Nosemosis negatively affects honeybee survival: experimental and meta-analytic evidence

**DOI:** 10.1101/2024.02.27.581909

**Authors:** M. Ostap-Chec, J. Cait, R. W. Scott, A. Arct, D. Moroń, M. Rapacz, K. Miler

**Author notes:** Corresponding authors: Monika Ostap-Chec, Krzysztof Miler. **Author’s contribution** Monika Ostap-Chec: Investigation, Funding acquisition, Writing – Original Draft, Writing – Review & Editing. Jessica Cait: Formal Analysis, Resources, Visualization, Writing – Review & Editing. R. Wilder Scott: Formal Analysis, Resources. Aneta Arct: Investigation, Writing – Review & Editing. Dawid Moroń and Marcin Rapacz: Resources, Writing – Review & Editing. Krzysztof Miler: Conceptualization, Methodology, Investigation, Formal analysis, Data Curation, Visualization, Resources, Project administration, Writing – Review & Editing.

## Abstract

Nosemosis, caused by microsporidian parasites of the genus *Nosema*, is considered a significant health concern for insect pollinators, including the economically important honeybee (*Apis mellifera*). Despite its acknowledged importance, the impact of this disease on honeybee survivorship remains unclear. Here, a standard laboratory cage trial was used to compare mortality rates between healthy and *Nosema*-infected honeybees. Additionally, a systematic review and meta-analysis of existing literature were conducted to explore how nosemosis contributes to increased mortality in honeybees tested under standard conditions. The review and meta-analysis included 50 studies that reported relevant experiments involving healthy and *Nosema*-infected individuals. Studies lacking survivorship curves or information on potential moderators, such as spore inoculation dose, age of inoculated bees, or factors that may impact energy expenditure, were excluded. Both the experimental results and meta-analysis revealed a consistent, robust effect of infection, indicating a threefold increase in mortality among the infected group of honeybee workers (hazard ratio for infected individuals = 3.16 [1.97, 5.07] and 2.99 [2.36, 3.79] in the experiment and meta-analysis, respectively). However, the meta-analysis also indicated high heterogeneity in the effect magnitude, which was not explained by our moderators. Furthermore, there was a serious risk of bias within studies and potential publication bias across studies. The findings underscore knowledge gaps in the literature. It is stressed that laboratory cage trials should be viewed as an initial step in evaluating the impact of *Nosema* on mortality and that complementary field and apiary studies are essential for identifying effective treatments to preserve honeybee populations.

## Introduction

Numerous insects in most climatic zones play important roles in pollination (Garibaldi *et al*. 2013; Klein *et al*. 2007). Among these, the honeybee (*Apis mellifera*) can be considered a key pollinator for many crops and wild plants due to the high efficiency of its workers in collecting and transferring pollen between flowers, which is essential for fertilization and subsequent fruit and seed production (Abrol 2012). The significance of honeybees is also underscored by their large colonies that enable the recruitment of high numbers of workers for pollination (Abrol 2012). This makes honeybees particularly valuable for commercial agriculture, where large-scale pollination is often key to ensuring high yields and the quality of crops (Aizen and Harder 2009; Morse and Calderone 2000). However, within the agricultural context, honeybees, and consequently beekeepers, face a wide range of significant issues (Genersch 2010). These encompass challenges such as exposure to pesticides and chemicals, along with susceptibility to diseases and parasites (Goulson *et al*. 2015; Potts *et al*. 2010).

Nosemosis is one of the most prevalent and widespread diseases in honeybees, often regarded as a significant threat to their health and well-being (Hristov *et al*. 2020; Moritz *et al*. 2010). This disease is caused by microsporidian parasites of the genus *Nosema*, which includes two species that infect honeybees: *N. apis* and *N. ceranae*. Although *Nosema* was recently reclassified as *Vairimorpha* (Tokarev *et al*. 2020), this revision has faced criticism (Bartolomé *et al*. 2024). To maintain clarity and consistency with the existing literature, we continue to refer to the genus as *Nosema*. Both parasites complete their life cycle within the honeybee midgut cells. Upon ingestion, spores reach the midgut, germinate, and inject their contents into host cells. Following phagocytosis, the infected cells are destroyed and release new spores (Gisder *et al*. 2011; Huang and Solter 2013), which ultimately result in gut lesions (reviewed in Goblirsch 2018). Consequently, affected bees become weakened and lethargic, which compromises the colony’s foraging capacity (Koch *et al*. 2017). The spores can infect other digestive tract cells or be excreted, contaminating nesting environments and floral resources. The fecal-oral transmission facilitates the easy and rapid spread of these parasites to new habitats and hosts (Fürst *et al*. 2014). Therefore, while *N. apis* was originally confined to Europe and North America, and *N. ceranae* to South East Asia, both species are now distributed worldwide (Chen *et al*. 2008a; Grupe and Quandt 2020; Paxton *et al*. 2007). The exacerbation of this spread can be at least in part attributed to the global trade of honeybee colonies and related products (Higes *et al*. 2008b; Mutinelli 2011).

Among the two agents responsible for nosemosis, *N. ceranae* has garnered significantly more attention compared to *N. apis*. *N. ceranae* was previously known as a parasite of *Apis ceranae* (Fries *et al*. 1996) and *A. mellifera* (Klee *et al*. 2007), but it has recently been confirmed to also infect other bee species (Grupe and Quandt 2020; Martín-Hernández *et al*. 2018). However, its infectivity varies depending on geographical origin and host species (Chaimanee *et al*. 2013; Müller et al. 2019; Porrini *et al*. 2020; van der Steen et al. 2022). *N. ceranae* appears to have higher biotic potential at different temperatures compared with *N. apis* (Martín-Hernández *et al*. 2009). The spread of *N. ceranae* is often associated with worker and colony mortality, including colony collapse syndrome (e.g., in Spain, (Botías *et al*. 2013; Higes *et al*. 2008a, 2009, 2010), but conflicting research suggests that nosemosis may not significantly contribute to beekeepers’ losses (e.g., (Cox-Foster *et al*. 2007; Fernández *et al*. 2012; Gisder *et al*. 2010; Kielmanowicz *et al*. 2015; Pohorecka *et al*. 2014; Schüler *et al*. 2023). Thus, the precise impact of *N. ceranae* infection on honeybee mortality remains unclear and requires further investigation.

Laboratory cage trials, a widely used standard method, involve the confinement of control and *Nosema*-infected honeybees in cages and strictly controlled conditions (Fries *et al*. 2013). Despite the frequent use of this method, primarily in the search for an effective treatment against the parasite (e.g., Borges *et al*. 2020; Chaimanee *et al*. 2021; Naree *et al*. 2021b; Van Den Heever *et al*. 2016), it has never been formally validated in the context of *Nosema* infection. Fries et al. (2013) compiled a list of commonly used methods, but their real-world applicability has never been thoroughly investigated. The described cage trial methods are general and leave researchers with a broad range of choices regarding specific conditions. Consequently, the trials are often poorly standardized across studies, leading to highly varied infection effects. For example, concerning mortality, some studies show a powerful effect of *N. ceranae* infection (e.g., Naree *et al*. 2021a; Van der Zee *et al*. 2014), while others indicate no discernible effect (e.g., Duguet *et al*. 2022; Hosaka *et al*. 2021; Huang *et al*. 2015; Retschnig *et al*. 2014a). Such discrepancies may stem from different experimental setups that involve, for example, various levels of artificial inoculation or ages of the hosts, both of which can affect nosemosis development (e.g., Berbeć *et al*. 2022; Jabal-Uriel *et al*. 2022). Additionally, considering that nosemosis operates through energy-related processes, such as hosts’ starvation or failed thermoregulation due to impaired digestive system function (Martín-Hernández *et al*. 2011; Mayack and Naug 2009; Vidau *et al*. 2014), assessing infection effects in honeybees maintained in laboratory cages with minimal energy expenditure, stable temperature, and *ad libitum* access to food might be suboptimal methodologically. Hence it is not clear how justified the use of this method is in *Nosema*-related research.

In this study, an experiment was conducted to compare mortality rates between healthy bees and those infected with *N. ceranae*, using a laboratory cage trial setup. Furthermore, a systematic review and meta-analysis of existing literature were performed to explore the impact of *N. ceranae* infection on mortality among honeybee workers in similar experimental setups. The primary objective was to assess the effect of nosemosis on honeybee survivorship to evaluate the suitability of mortality assessment in caged honeybees as a method. It was hypothesized that, generally, *N. ceranae* infection would lead to a significant decrease in honeybee survivorship. However, it was also hypothesized that the magnitude of this effect would depend on mediating factors that often vary between studies, such as the spore dosage used for inoculation or the age of the experimentally inoculated honeybees. Incubation temperature and food supplementation were included as additional factors that may impact energy expenditure and serve as potential mediators of the effect of infection on mortality. All these factors are likely to influence the development and course of nosemosis (Goblirsch 2018).

## Materials and methods

### Survivorship experiment

#### Preparation of spores for infection and their genetic identification

The spore suspension for experimental infection was freshly prepared. Utilizing a stock population of infected honeybees, the digestive tract of several individuals was homogenized with a micro-pestle and suspended in distilled water. To isolate spores, the suspension underwent centrifugation (Frontier 5306, Ohaus, Switzerland) for 5 minutes at 2,000 G, repeated three times, with each round involving the replacement of the supernatant with fresh distilled water. Subsequently, the supernatant was substituted with a 1M sucrose solution, and the spore concentration was assessed using a Bürker hemocytometer under a Leica DMLB light microscope equipped with phase contrast (PCM) and a digital camera. Achieving a final concentration of 100,000 spores per 10 µL involved appropriate dilution of the infection solution with 1M sucrose solution. This dose was subsequently used for individual infection in the experiment (a typical dose used to ensure infection Fries *et al*. 2013). To verify the identity of *N. ceranae* spores, PCR was employed following the protocol outlined by Berbeć et al. (Berbeć *et al*. 2022). Briefly, 50 µL of the spore suspension was incubated in TNES buffer (100 mM Tris-HCl pH 8.0, 5 mM EDTA, 0.3% SDS, 200 mM NaCl) with 8 µL of proteinase K (10 mg/ml) for 2 hours at 56°C with shaking. Following centrifugation, DNA from the supernatant was precipitated by adding an equal volume of 100% isopropanol, washed twice in 70% ethanol, and resuspended in 50 µL of nuclease-free water. PCR amplification, using species-specific primers complementary to the rRNA genes of *Nosema* species and a PCR Mix Plus kit containing PCR anti-inhibitors (A&A Biotechnology), was carried out under the following conditions: 94°C for 3 minutes for initial denaturation and 35 cycles (94°C for 30 seconds, 52°C for 30 seconds, and 72°C for 30 seconds). Primers were employed following Chen et al. (Chen *et al*. 2008b). The resulting amplification products were analyzed through gel electrophoresis to confirm the exclusive presence of *N. ceranae*.

#### Experimental procedure

We obtained newly emerged worker honeybees (*Apis mellifera carnica*) from two unrelated and queenright colonies with naturally inseminated queens. The selected colonies were in overall good condition, having undergone treatment with oxalic acid against *Varroa destructor* in early spring. Additionally, before the experiment, they were confirmed to be *Nosema*-free through spore count assessment in several randomly collected foragers using hemocytometry. For the experiment, we obtained worker bees by selecting a single bee-free frame with capped brood from each colony and placing it in an incubator (KB53, Binder, Germany) at 32 °C overnight. The emerged bees were used for individual feeding in the laboratory. About 120 individuals were placed on Petri dishes (30 bees per group and colony) and left for approximately 1 hour to increase their feeding motivation. Following this, a droplet of food was provided to each bee. In the experiment, individuals in the infected groups (one from each colony) received a 10 µl drop of a 1M sucrose solution containing 100,000 spores of *N. ceranae*, while individuals in the control groups (another from each colony) received a 10 µl drop of a 1M sucrose solution without spores. The bees were monitored for 3 hours, and any bee that consumed the food was immediately transferred to the appropriate cage. Bees that failed to consume the provided food were excluded. Ultimately, we achieved a final count of 20 bees in each cage. The cages were provided with *ad libitum* water and gravity feeders with 1M sucrose solution and were placed in an incubator (KB400, Binder, Germany) at 34 °C. Water and food were renewed every morning in each cage. The mortality assessment lasted for 38 days, continuing until the death of all individuals, and was performed blind (cages were coded). We refrained from counting weakened or lethargic individuals as dead, leaving them in their cages. All dead individuals were frozen and later analyzed to confirm their infection status (control vs. infected).

#### Analysis of experimental samples

The level of infection in honeybees after the survivorship experiment was examined using hemocytometry. The digestive tract of each frozen honeybee was homogenized using micro-pestle in 300 μL distilled water. The spores were counted in a hemocytometer, analogically to the preparation of spores for infection. The contents of 5 small squares (volume: 0.00125 μL) were counted. If the number of spores counted per sample was less than 10, the contents of 5 large squares (volume: 0.02 μL) were counted. To determine the total number of spores per individual we used the following formula: number of spores per individual = number of spores per sample × 300 μL / total solution volume of sample. We analyzed 40 control and 39 infected honeybees in total (1 infected sample was misplaced).

#### Statistical analysis

To analyze the survivorship data, we used the Cox mixed-effect regression with a fixed factor of the group and random factor of the colony in R (coxme, survival, and survminer packages) and visualized the results using Kaplan-Meier curves (Kassambara *et al*. 2021; Therneau 2022, 2023). We chose Cox mixed-effect regression for our survival analysis to account for random effects and estimate hazard ratios in the control and infected individuals.

### Systematic review and meta-analysis

We followed the Preferred Reporting Items for Systematic Reviews and Meta-Analyses (PRISMA) guidelines (Moher *et al*. 2015). The review and its protocol have not been pre-registered.

#### Eligibility criteria

We first prepared a list of eligibility criteria for studies to be included in the systematic review and meta-analysis: (i) in English, (ii) with the use of workers of the honeybee (*Apis mellifera*), (iii) comprising a laboratory experiment with a treatment group, i.e. workers exposed to *N. ceranae* spores, and a control group, i.e. workers exposed to no *Nosema* treatment, and (iv) reporting a measure of mortality in the form of a survivorship curve.

#### Data sources and search

We completed an electronic search of documents on October 31, 2022. For this, we used Web of Science™ and the Scopus databases. Our search used the phrase “nosema mortality apis” in the topic, as well as a forward search (i.e. documents citing one or more works on the list) refined by the same phrase in the topic. References were de-duplicated using Mendeley. The screening was conducted in two phases. First, we screened the documents for potential inclusion based on their titles and abstracts. Second, we retrieved the full texts of potentially eligible documents for further reading. This action was done independently by two investigators and the full texts retrieved by just one or both were used.

#### Extraction of study details and study exclusion

From the selected studies, we extracted the following types of data: (i) inoculation dose (spores per individual), (ii) age of the inoculated workers, (iii) food provided to the workers (plain sucrose or with supplements), (iv) the number of individuals per cage as well as the number of replicates and the resulting group size (N), (v) days of mortality assessment (i.e. duration of the survivorship analysis), and (vi) incubation temperature. Further, we noted (vii) the inoculation method (individual or collective) as a factor likely to compromise the accuracy of the above-mentioned data type (i) and, as such, a potential methodological mediator of the effect of infection on mortality. This action was completed by one investigator. Studies were excluded if (i) survivorship figures were unreadable, (ii) infection dose or method was not specified, (iii) group size was not reported, (iv) *Nosema* spores used were of mixed species, (v) worker age was unknown. There was also one case in which the survivorship was reported for only 24 hours and another case in which the results were already reported in a different publication (duplicate results). Both these works were excluded. For the full list of references containing the studies excluded during the second phase of screening see Appendix A.

Among the included studies, some did not specify the number of replicates (4 studies) or incubation temperature (1 study). In these cases, we noted the number of replicates as 1 and the temperature as 33 °C (the median from the other studies). Further, some works used multiple *Nosema* populations with different geographic origins (1 study) and multiple inoculation doses (3 studies), but with a single control group of workers. Thus, a single infected group of workers had to be chosen. In the case of multiple *Nosema* populations, we selected the group infected by spores from a population corresponding geographically to the honeybee population. In the case of multiple inoculation doses, we selected the group with the lowest inoculation dose as studies that utilize low doses are rare.

#### Risk of bias

The risk of bias was assessed by two investigators independently, with disagreements resolved by consensus, using a modified SYRCLE tool (Hooijmans et al. 2014) and visualized with a summary plot in R using the robvis package (McGuinness 2019). We used the following assessment categories: (1) whether baseline characteristics of animals allocated to the control and infected groups were likely similar (low bias if were similar, high bias if were not similar, unclear if there were differentiating factors of unknown effect), (2) whether housing conditions in the incubator were randomized (low if cage positions were switched, high if cages were stationary, unclear if randomization was not mentioned), (3) whether investigators assessing mortality were blind to the group identity of animals (low bias if were blind, high bias if were not blind, unclear if blinding was not mentioned), (4) whether the contamination was measured and reported (low bias if was measured and reported, high bias if was not measured or not reported, unclear if was not mentioned), (5) whether there was any commercial funding for the study (low bias if none, high bias if funded by industry, unclear if was not mentioned). These assessment categories had the highest relevance in the context of our systematic review and meta-analysis.

#### Data extraction for survivorship

To assess worker mortality in the selected studies, we computed hazard ratios (HRs) by reconstructing Kaplan-Meier curves from figures included in the papers using a Web Plot Digitizer (Rohatgi 2022). This action was performed first by one investigator, and then checked for completeness and accuracy by another investigator, with corrections applied if needed by consensus. Curve construction, HRs, and confidence interval calculations were performed via methods and R script from Guyot et al. (Guyot *et al*. 2012).

#### Meta-analysis

Analyses were conducted in R using the random-effects meta-analysis function (rma) in the metafor package (Viechtbauer 2010). Data are reported such that increased death in infected groups resulted in HR > 1.

#### Publication bias

Publication bias, the failure to publish null results from small studies, was assessed visually by examining the degree of effect asymmetry in the funnel plot (standard error plotted against HRs, Appendix B) and was evaluated statistically via the rank correlation test.

#### Exploration of data heterogeneity

Heterogeneity was assessed using the Q and I^2^ statistics. To explore potential causes of data heterogeneity, we conducted meta-regressions. These regressions included inoculation dose, inoculation age of workers, and incubation temperature as continuous moderators, and food supplementation (sucrose or supplement) and inoculation method (individual or collective) as categorical moderators in the random-effects model.

#### Quality of evidence

The strength of the body of evidence was assessed by one investigator using the Grading of Recommendations, Assessment, Development, and Evaluation (GRADE) guidelines (Guyatt *et al*. 2011). The quality of evidence was based on the study design and decreased for risk of bias, publication bias, inconsistencies, indirectness, or imprecision in the included studies as well as increased for the size of the effect or dose-response relationship.

## Results

### Survivorship experiment

Cox regression showed a significant effect of the infection (control vs. infected group of workers, z = 4.77, p < 0.001). The hazard ratio for the group infected with *N. ceranae* spores was 3.16 [1.97, 5.07] (Fig. 1), indicating that the risk of death in infected bees was three times higher than in healthy bees. The infected group had high infection levels (average number of spores per individual: 18,536,634, 39 individuals in total) with 8 individuals that had no detectable spores. In the control group, we found 5 *Nosema*-contaminated samples (range: 30,000-165,000 spores per individual, 40 individuals in total).

**Fig. 1.**
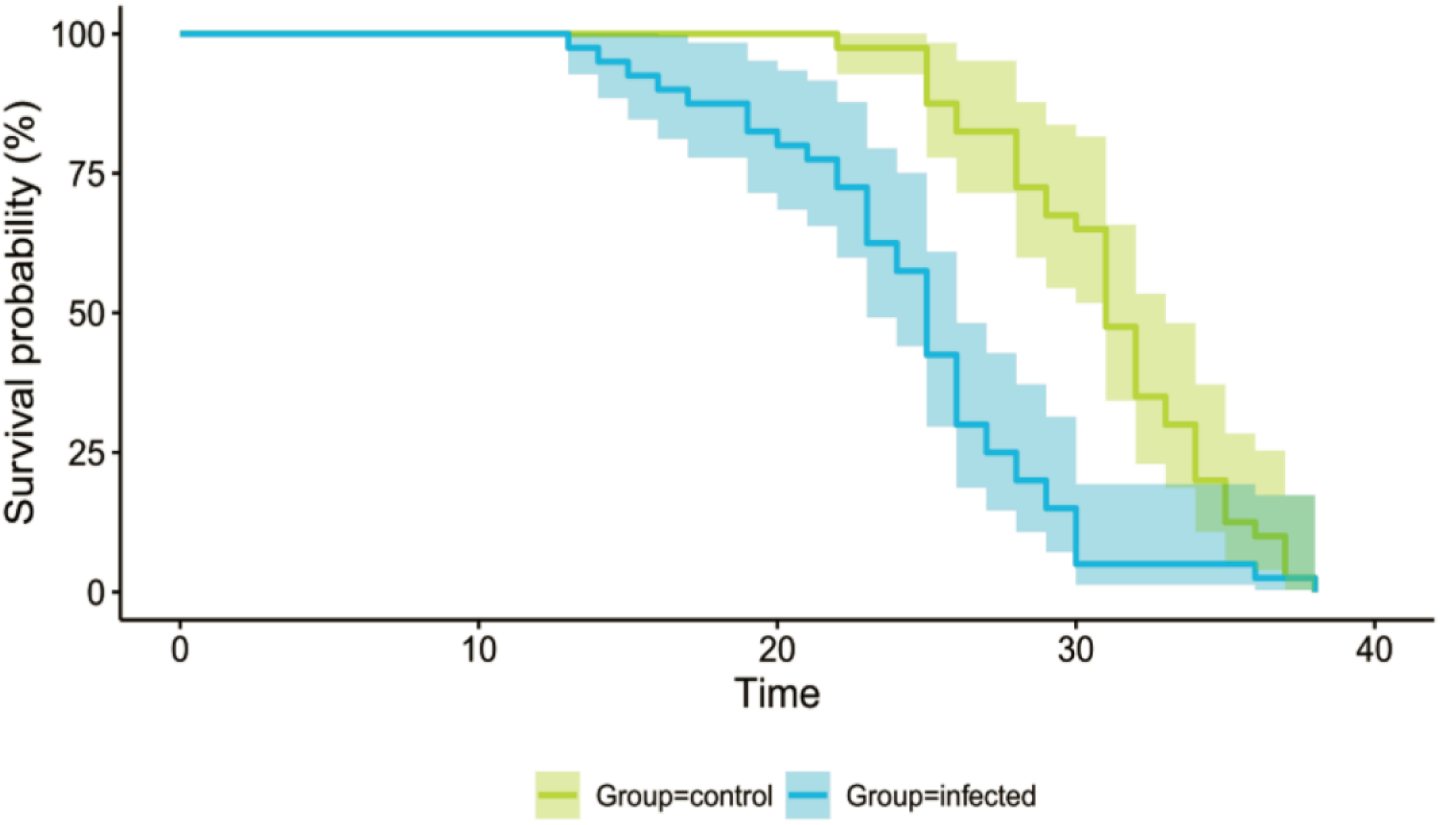
Kaplan-Meier survival curves for the control and infected groups of workers. The two groups differ significantly in survival. Shading indicates 95 % confidence intervals.

### Systematic review and meta-analysis

#### Study characteristics

Our search yielded 1599 documents in total (273 from Web of Science™ and 1326 from Scopus). After de-duplication (n = 314 duplicates) and removal of patents (n = 1) 1284 documents were screened for potential inclusion based on their titles and abstracts. This action yielded a total of 123 publications. Upon reading the full texts of the selected 123 publications, we were able to identify 52 eligible studies. During data extraction for survivorship, we excluded 1 additional article (containing 2 studies that used separate groups of animals) because no mortality occurred in the control groups for over 2 weeks, which made resulting hazard ratios improbably large and thus unusable. By this action, the number of studies included in the meta-analysis decreased to 50 from 44 publications (Fig. 2).

**Fig. 2.**
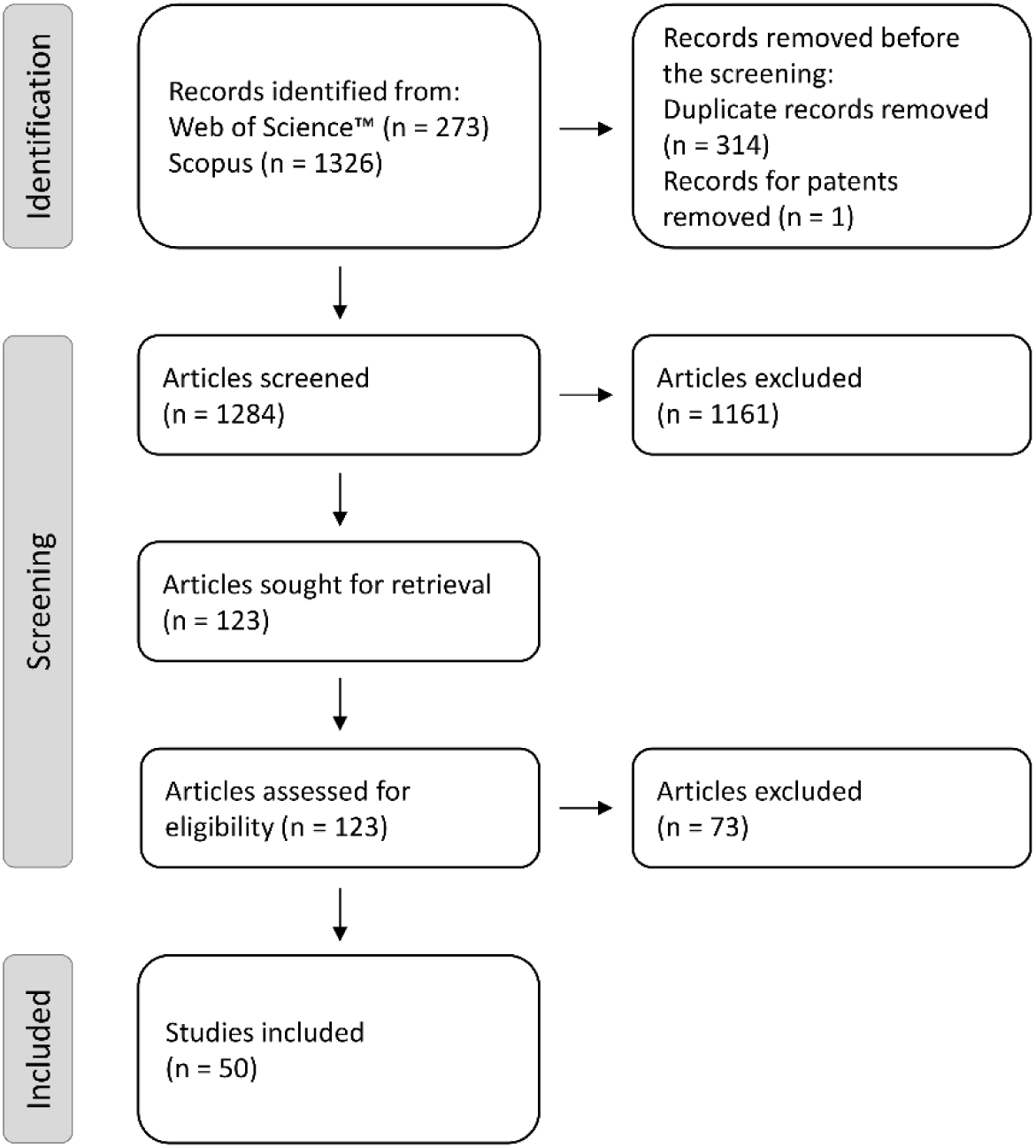
Preferred Reporting Items for Systematic Reviews and Meta-Analyses (PRISMA) flow diagram.

Among the included studies (Table 1), the majority (70 %) used a plain sucrose diet, while 30 % of the studies included diet supplementation. Most of these supplements involved a protein source, with 9 studies using commercially available protein patties such as Provita’Bee, Pro Bee, or MegaBee, 5 studies using handmade pollen paste or beebread and 1 study using Apifonda powdered sugar. Similarly, most studies implemented individual (82 %), not collective spore feeding method (18 %). Freshly emerged, one-day-old workers were most often used (in 38 % of studies) and an inoculation dose of 100,000 spores per individual was most often implemented (in 42 % of studies). In the majority of works included in the meta-analysis, the mortality assessment lasted for longer than 2 weeks (70 %), and the workers undergoing assessment were most frequently kept at 33 or 34 °C (in 58 % of studies).

**Table 1.** Overview of studies included in the systematic review and meta-analysis. The table presents information on studies investigating worker honeybee mortality in control and *Nosema*-infected individuals. Each study is catalogued with details including reference, the type of food provided (plain sucrose or supplemented), worker age at spore inoculation, method of inoculation (individual or collective feeding), estimated spore dose per individual, group sizes with replicates, total number of tested individuals (N), duration of mortality assessment, and incubation temperatures.

| Study | Reference | Food | Inoculation age (days) | Inoculation method | Inoculation dose | Group size | Number of replicates | N | Mortality assessment (days) | Temperature |
| --- | --- | --- | --- | --- | --- | --- | --- | --- | --- | --- |
| 1 | Duguet et al. 2022 | Plain sucrose | 3 | Individual | 120,000 | 20 | 4 | 80 | 14 | 34 |
| 2 | Berbeć et al. 2022 | Plain sucrose | 2 | Collective | 10,000 | 150 | 4 | 600 | 7 | 30 |
| 3 |  | Plain sucrose | 10 | Collective | 10,000 | 150 | 4 | 600 | 7 | 30 |
| 4 | Balbuena et al. 2023 | Plain sucrose | 2 | Collective | 100,000 | 70 | 3 | 210 | 30 | 30 |
| 5 | Zhang et al. 2021 | Plain sucrose | 5 | Individual | 100,000 | 45 | 3 | 135 | 20 | 34 |
| 6 | Özgör 2021 | Plain sucrose | 1 | Individual | 1,000,000 | 30 | 1 | 30 | 12 | 34 |
| 7 | Naree et al. 2021b | Supplement | 1 | Individual | 100,000 | 50 | 3 | 150 | 30 | 34 |
| 8 | Liu et al. 2021 | Plain sucrose | 10 | Individual | 100,000 | 20 | 3 | 60 | 10 | 35 |
| 9 | Hosaka et al. 2021 | Plain sucrose | 1 | Individual | 140 | 30 | 3 | 90 | 21 | 33 |
| 10 | Chaimanee et al. 2021 | Plain sucrose | 3 | Individual | 100,000 | 30 | 3 | 90 | 10 | 34 |
| 11 | Borges et al. 2021 | Plain sucrose | 1 | Individual | 50,000 | 40 | 1 | 40 | 54 | 30 |
| 12 | Almasri et al. 2021 | Plain sucrose | 1 | Individual | 100,000 | 30 | 7 | 210 | 21 | 33 |
| 13 | Straub et al. 2020 | Supplement | 1 | Collective | 10,000 | 20 | 22 | 440 | 14 | 34 |
| 14 | Porrini et al. 2020 | Supplement | 2 | Individual | 1,000,000 | 50 | 5 | 250 | 13 | 28 |
| 15 | Paris et al. 2020 | Plain sucrose | 2 | Collective | 100,000 | 50 | 3 | 150 | 22 | 33 |
| 16 | Mura et al. 2020 | Plain sucrose | 1 | Individual | 100,000 | 21 | 3 | 63 | 30 | 31 |
| 17 | Kim et al. 2020 | Plain sucrose | 4 | Individual | 50,000 | 20 | 3 | 60 | 15 | 34 |
| 18 |  | Plain sucrose | 4 | Individual | 50,000 | 20 | 3 | 60 | 15 | 34 |
| 19 |  | Plain sucrose | 4 | Individual | 50,000 | 20 | 3 | 60 | 15 | 34 |
| 20 |  | Plain sucrose | 4 | Individual | 50,000 | 20 | 3 | 60 | 15 | 34 |
| 21 | Borges et al. 2020 | Plain sucrose | 1 | Individual | 50,000 | 40 | 3 | 120 | 54 | 33 |
| 22 | Bell et al. 2020 | Plain sucrose | 1 | Individual | 40,000 | 123 | 1 | 123 | 9 | 36 |
| 23 | Arismendi et al. 2020 | Supplement | 2 | Individual | 100,000 | 75 | 4 | 300 | 20 | 30 |
| 24 | Sinpoo et al. 2018 | Plain sucrose | 5 | Individual | 100,000 | 30 | 3 | 90 | 14 | 34 |
| 25 | Ptaszyńska et al. 2018 | Plain sucrose | 3 | Collective | 125,000 | 40 | 10 | 400 | 16 | 35 |
| 26 | Panek et al. 2018 | Supplement | 1 | Individual | 50,000 | 130 | 4 | 520 | 21 | 33 |
| 27 | Li et al. 2018 | Supplement | 1 | Individual | 100,000 | 40 | 4 | 160 | 21 | 34 |
| 28 | Arredondo et al. 2018 | Plain sucrose | 3 | Collective | 500,000 | 50 | 3 | 150 | 7 | 35 |
| 29 | Tritschler et al. 2017 | Plain sucrose | 1 | Collective | 100,000 | 50 | 6 | 300 | 14 | 35 |
| 30 | Paris et al. 2017 | Supplement | 6 | Individual | 100,000 | 45 | 24 | 1080 | 22 | 33 |
| 31 | Li et al. 2017 | Plain sucrose | 2 | Individual | 100,000 | 40 | 6 | 240 | 26 | 34 |
| 32 | Ptaszyńska et al. 2016 | Plain sucrose | 3 | Individual | 32,000 | 40 | 6 | 240 | 21 | 30 |
| 33 | Natsopoulou et al. 2016 | Plain sucrose | 4 | Individual | 100,000 | 18 | 5 | 90 | 42 | 30 |
| 34 | Higes et al. 2016 | Plain sucrose | 5 | Individual | 50,000 | 25 | 3 | 75 | 30 | 27 |
| 35 | Milbrath et al. 2015 | Plain sucrose | 1 | Individual | 30,000 | 100 | 3 | 300 | 30 | 30 |
| 36 | Huang et al. 2015 | Supplement | 5 | Individual | 100,000 | 30 | 3 | 90 | 50 | 30 |
| 37 | Doublet et al. 2015 | Plain sucrose | 2 | Individual | 100,000 | 30 | 3 | 90 | 13 | 30 |
| 38 |  | Plain sucrose | 2 | Individual | 100,000 | 30 | 4 | 120 | 25 | 30 |
| 39 | Williams et al. 2014 | Plain sucrose | 2 | Individual | 35,000 | 20 | 3 | 60 | 30 | 33 |
| 40 | Van der Zee et al. 2014 | Plain sucrose | 5 | Individual | 50,000 | 25 | 4 | 100 | 25 | 34 |
| 41 | Retschnig et al. 2014b | Supplement | 2 | Individual | 100,000 | 20 | 5 | 100 | 14 | 34 |
| 42 | Retschnig et al. 2014a | Plain sucrose | 1 | Collective | 100,000 | 20 | 4 | 80 | 14 | 30 |
| 43 | Aufauvre et al. 2014 | Supplement | 1 | Individual | 125,000 | 165 | 1 | 165 | 25 | 33 |
| 44 | Milbrath et al. 2013 | Plain sucrose | 5 | Individual | 125,000 | 20 | 2 | 40 | 32 | 33 |
| 45 | Goblirsch et al. 2013 | Supplement | 1 | Individual | 10,000 | 30 | 3 | 90 | 28 | 28 |
| 46 | Dussaubat et al. 2013 | Plain sucrose | 5 | Individual | 40,000 | 30 | 3 | 90 | 19 | 33 |
| 47 | Di Pasquale et al. 2013 | Supplement | 1 | Individual | 100,000 | 70 | 1 | 70 | 50 | 34 |
| 48 | Aufauvre et al. 2012 | Supplement | 1 | Individual | 125,000 | 50 | 3 | 150 | 22 | 33 |
| 49 |  | Supplement | 1 | Individual | 125,000 | 50 | 3 | 150 | 22 | 33 |
| 50 | Vidau et al. 2011 | Supplement | 5 | Individual | 125,000 | 50 | 3 | 150 | 20 | 35 |

#### Risk of bias

Overall, the risk of bias was serious (Fig. 3). Baseline characteristics of animals participating in the studies were mostly similar between groups, except for a few studies in which the control and infected groups were kept in separate housing incubators. Randomization of the position of cages within the housing incubator was applied in only one study, thanks to which it avoided possible effects related to the position of a cage in the incubator. Notably, not a single study mentioned any form of blinding applied during the mortality assessment. Considering that only one study listed specific criteria for counting an individual as dead, this made the omission of blinding quite conspicuous and created space for investigator bias during mortality assessment. Several studies failed to control the contamination of the control groups and infection levels in the artificially infected groups. There was no commercial funding in any of the studies.

**Fig. 3.**
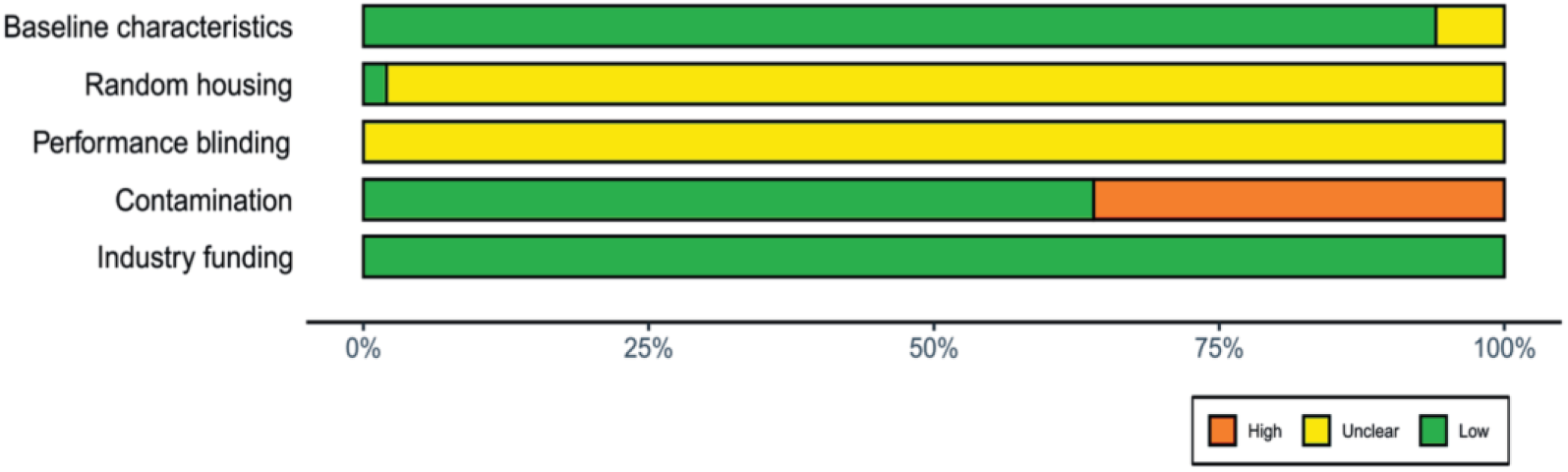
Summary plot of the results from the Systematic Review Centre for Laboratory animal Experimentation (SYRCLE) risk of bias tool. Green indicates a low risk of bias, yellow is unclear, and red indicates a high risk of bias.

#### Meta-analysis

A random-effects meta-analysis of the hazard ratios (HRs) showed a significant effect of infection with *N. ceranae* spores (HR = 2.99 [2.36, 3.79]) (Fig. 4), indicating that infected bees had nearly three times higher risk of death compared to healthy bees. There was high heterogeneity (Q = 647.32, df = 49, p < 0.0001; I² = 94.44%).

**Fig. 4.**
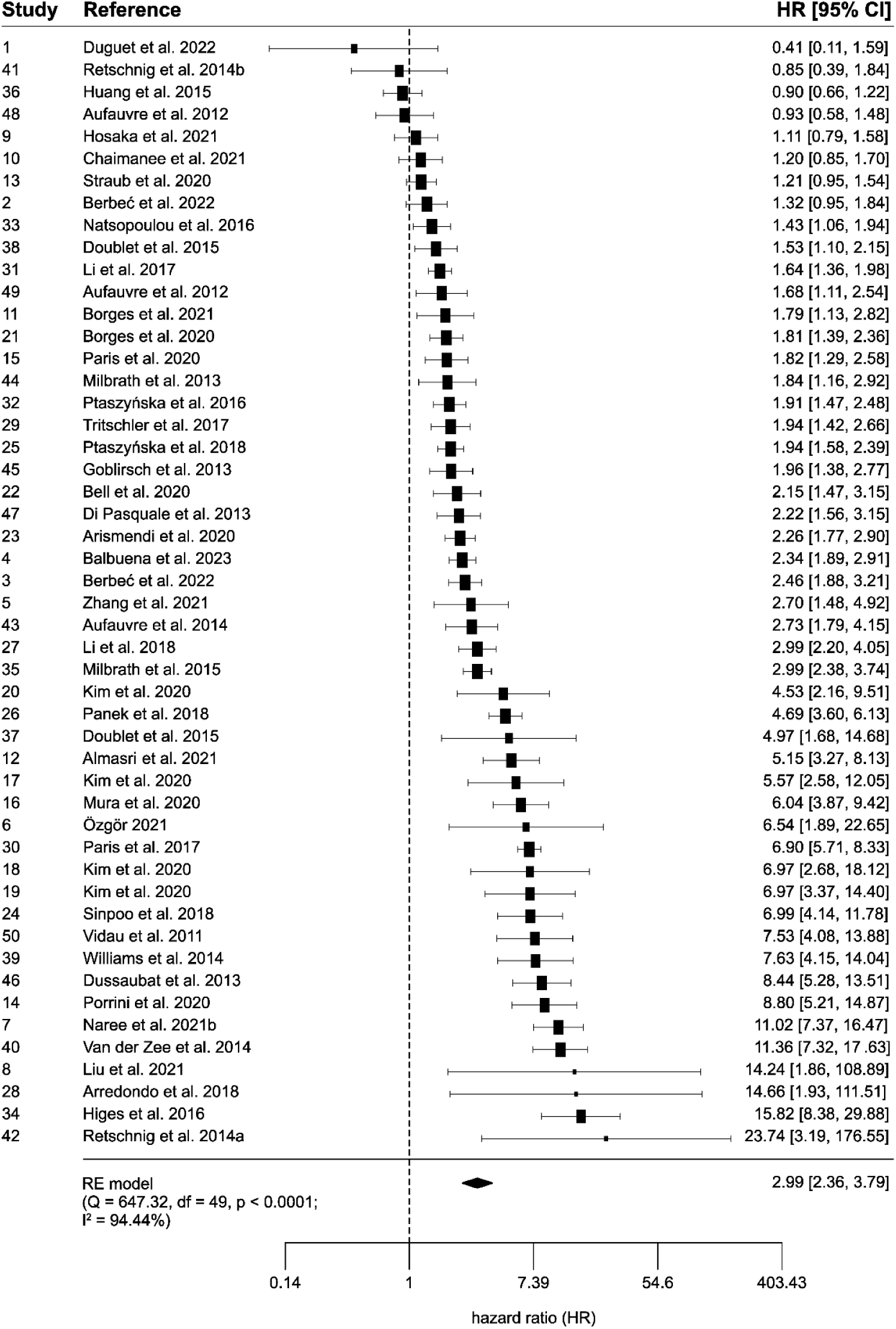
Forest plot of the random-effects meta-analysis of hazard ratios (HRs) and 95 % confidence intervals. Diamond demonstrates the overall estimate (with the width reflecting the 95 % CI). Black squares indicate the study HRs and their sizes indicate the weight of the study in the meta-analysis. The dashed line indicates no difference between groups (HR = 1). The Q and I^2^ statistics are tests of heterogeneity.

#### Publication bias

Visual inspection of the funnel plot and the rank correlation test indicated the presence of publication bias (tau = 0.223, p = 0.022) (Appendix B).

#### Exploration of data heterogeneity

Included moderators did not significantly explain HR magnitudes (F_5,44_ = 1.812, p = 0.130, R^2^ = 10.92 %). A trend emerged demonstrating that HRs increased with increasing inoculation doses (F_1,44_ = 3.650, p = 0.063). Another trend indicated a positive relationship between the increased age of workers at inoculation and larger HRs (F_1,44_ = 3.800, p = 0.058).

#### Quality of evidence

All studies included in the systematic review and meta-analysis were considered randomized trials. The overall quality of evidence was high (Table 2). Even with a serious risk of bias, likely publication bias, and serious inconsistencies indicated by high heterogeneity, the included studies had no serious indirectness or imprecision. Moreover, the detected effect was large and there was a trend towards dose response.

**Table 2.** Grading of Recommendations, Assessment, Development, and Evaluations (GRADE) assessment of confidence of cumulative evidence.

| Number of studies | Overall risk of bias | Inconsistencies | Indirectness | Imprecision | Publication bias | Effect | Dose response | Quality |
| --- | --- | --- | --- | --- | --- | --- | --- | --- |
| 50 | Serious | Serious | Not serious | Not serious | Likely | HR = 2.99<br>[2.36, 3.79]<br>(large) | F<sub>1,44</sub> = 3.650,<br>p = 0.063 | High |

## Discussion

Both our experiment and meta-analysis indicated a consistent and strong impact of infection, with mortality increasing about threefold in the group of honeybees infected with *N. ceranae* compared to uninfected individuals. Importantly, we found no evidence of the incubation temperature, food supplementation, or inoculation method influencing the magnitude of the mortality effect. While trends suggested higher infection effects in studies where honeybees were infected with larger doses of spores or in studies involving older hosts, the moderators collectively accounted for only 10.92 % of the large heterogeneity in HRs (I^2^ = 94.44 %). The results from our experiment closely aligned with the findings of the meta-analysis. Thus, our experiment can serve as a valuable reference for standardizing research methodologies in laboratory cage trials.

The lack of strong evidence regarding the significance of any of the included moderators was surprising. Future studies must account for high heterogeneity in the effect of *Nosema* infection on honeybees. Given the type of studies incorporated into our systematic review and meta-analysis (Table 1), we strongly recommend future research to address the impact of infection on mortality in older individuals, including those considerably older than a few days. Most of the research included in this analysis focuses on one-day-old bees, which is somewhat justified since newly emerged bees are more susceptible to *Nosema* infection compared to older individuals (Urbieta-Magro *et al*. 2019). Additionally, even in healthy colonies, trace amounts of *Nosema* spores are detectable, suggesting that the in-hive environment may be the first where bees encounter this pathogen. However, several studies have failed to detect spores in one-day-old bees while consistently identifying *N. ceranae* spores in older bees (Jack *et al*. 2016; Smart and Sheppard 2012). This indicates that the external environment is another source of *N. ceranae* infection. Foraging for food likely exposes bees to spores, increasing their risk of infection. Our research suggests a trend for a positive correlation between the age of workers at the time of inoculation and the risk of mortality. Therefore, it is crucial to investigate the effects of repeated exposure to low doses of spores in older bees, particularly foragers. This experimental setup would provide a more accurate approximation of real-world scenarios, addressing a significant gap in our understanding of *Nosema* infection dynamics. Moreover, this scenario could help explain the extremely high spore infection levels, often reaching multi-million counts, observed in the oldest bees (Jabal-Uriel *et al*. 2022; Li *et al*. 2017; Smart and Sheppard 2012). Additionally, another important issue for future research would be the geographic patterns. Across continents, the majority of the studies included here were based in Europe, which may disproportionately reflect conditions, practices, or bee populations specific to this region. Geographic bias needs to be addressed to enable global analyses and comparisons between regions, especially considering that experiments devoted to geographic differences in *Nosema* virulence are scarce (e.g., Genersch 2010; Dussaubat *et al*. 2013; Van der Zee *et al*. 2014). Other potentially important factors that require further study include, for example, the phenotype of the studied honeybees or the season (time of year).

Our findings indicate that *N. ceranae* is a serious threat to honeybees in terms of mortality. This study strongly supports the idea that nosemosis can significantly contribute to honeybee colony losses. However, it’s crucial to treat cage trials as an initial step in assessing infection effects and possible treatment measures. The subsequent step should involve conducting similar research under natural conditions within colonies, where bees experience not only individual immunity but also full social immunity. The value of this approach is confirmed when we compare the results of studies such as those by Li et al. (Li *et al*. 2018) and Lourenço et al. (Lourenço *et al*. 2021). In the former, artificially infected bees were examined in cages, while in the latter, artificially infected bees were introduced into colonies. The results showed a significantly lower spore load in bees living in colonies compared to those maintained in cages 12 days post-infection. This demonstrates the significant role that the colony environment and its components, including substances with high antibiotic activity like propolis (Mura *et al*. 2020; Simone-Finstrom *et al*. 2017), secondary plant metabolites in pollen and honey (Erler and Moritz 2016), or even queen presence (Huang *et al*. 2024), can play.

Equally important are the behavioral defense mechanisms of bees, such as avoiding trophallaxis among sick bees (Naug and Gibbs 2009) or consumption of honey with higher antibiotic activity by diseased bees (Gherman *et al*. 2014). In natural colony conditions, however, honeybees face various challenges, including exposure to pesticides and other chemical substances. The synergistic impact of these challenges, combined with *Nosema* infection, surely increases the mortality rate among bees and affects queen survival (Aufauvre *et al*. 2012, 2014; Dussaubat *et al*. 2016; Paris *et al*. 2018). Additionally, *Nosema*-infected bees experience higher levels of hunger, leading to energetic stress (Alaux *et al*. 2010; Mayack and Naug 2009). The expenditure of energy in the colony environment is considerably higher than in laboratory cages due to task performance. Therefore, not only should experimental methodologies be critically evaluated for ecological accuracy, but outcomes obtained in controlled settings need validation within the context of natural environments. This approach is essential to ensure that the results and conclusions capture the complexity of real-world effects.

Given that our results indicate a high risk that *Nosema* poses to honeybees, it is important to consider these findings in relation to other bee species. This is particularly concerning due to the ease with which *N. ceranae* can infect a variety of species, including bumblebees and solitary bees (Grupe and Quandt 2020; Martín-Hernández et al. 2018; Porrini et al. 2017). However, our findings should not be seen as an unequivocal threat to all bee species. For instance, studies on the solitary bee *Osmia bicornis* suggest that the impact of *Nosema* infection on mortality is minimal (Müller et al. 2019). In the case of bumblebees, although *N. ceranae* has been detected in various species (Plischuk et al. 2009; Grupe and Quandt 2020), recent evidence shows that spores pass through the digestive tract without leading to infection, suggesting no pathogen proliferation (van der Steen et al. 2022). Therefore, *N. ceranae* may not represent as severe a threat to solitary bees and bumblebees as it does to honeybees. Instead, these species may act as reservoir hosts for *N. ceranae* within pollinator networks.

Our study indicates that *N. ceranae* significantly influences the mortality of worker bees and, as such, likely poses a serious survival threat to the honeybee and the ecosystem services it provides (Papa *et al*. 2022). The search for the treatment of nosemosis is ongoing (Iorizzo *et al*. 2022). Significant attention is paid to the application of Good Beekeeping Practices as a form of prevention (Formato *et al*. 2022). However, while these methods yield some results, they still prove insufficient (Garrido *et al*. 2024; Holt and Grozinger 2016; Huang *et al*. 2013; Lang *et al*. 2023; Prouty *et al*. 2023). There is a pressing need for further research on *N. ceranae*, especially in the context of its effective control strategies that consider the microbiological safety of the hive products (i.e. pollen and honey).

## Acknowledgements

We thank Sławomir Jurczyk (University of Agriculture in Krakow, Poland) for technical assistance.

