## Appendix A for "Nosemosis negatively affects honeybee survival: experimental and meta-analytic evidence"

[1]Doctoral School of Exact and Natural Sciences, Jagiellonian University, Kraków, Poland; [2]Institute of Environmental Sciences, Faculty of Biology, Jagiellonian University, Kraków, Poland; [3]Department of Integrative Biology, College of Biological Science, University of Guelph, Guelph, ON, Canada; [4]Biological Sciences Platform, Holland Bone and Joint Program, Sunnybrook Research Institute; Department of Laboratory Medicine & Pathobiology, University of Toronto, ON, Canada; [5]Institute of Systematics and Evolution of Animals of the Polish Academy of Sciences, Kraków, Poland; [6]Department of Plant Physiology, Breeding and Seed Science, Faculty of Agriculture and Economics, University of Agriculture in Krakow, Poland

Corresponding authors:

Monika Ostap-Chec,

Krzysztof Miler,

References (excluded at the second phase of screening):


1. Al Naggar, Y., Baer, B., 2019. Consequences of a short time exposure to a sublethal dose of Flupyradifurone (Sivanto) pesticide early in life on survival and immunity in the honeybee (*Apis mellifera*). Sci. Rep. 9, 19753.
2. Alaux, C., Brunet, J., Dussaubat, C., Mondet, F., Tchamitchan, S., Cousin, M., Brillard, J., Baldy, A., Belzunces, L. P., Le Conte, Y., 2010. Interactions between *Nosema* microspores and a neonicotinoid weaken honeybees (*Apis mellifera*). Env. Microbiol. 12, 774–782.
3. Alberoni, D., Di Gioia, D., Baffoni, L., 2023. Alterations in the Microbiota of Caged Honeybees in the Presence of *Nosema ceranae* Infection and Related Changes in Functionality. Microbial Ecol. 86, 601–616.
4. Andrearczyk, S., Kadhim, M. J., Knaga, S., 2014. Influence of a probiotic on the mortality, sugar syrup ingestion and infection of honeybees with *Nosema* spp. under laboratory assessment. Med. Weter. 70, 762–765.
5. Baffoni, L., Gaggìa, F., Alberoni, D., Cabbri, R., Nanetti, A., Biavati, B., Di Gioia, D., 2016. Effect of dietary supplementation of *Bifidobacterium* and *Lactobacillus* strains in *Apis mellifera* L. against *Nosema ceranae*. Benef. Microbes 7, 45–51.
6. Bahreini, R., Currie, R. W., 2015. The influence of *Nosema* (Microspora: Nosematidae) infection on honey bee (Hymenoptera: Apidae) defense against *Varroa destructor* (Mesostigmata: Varroidae). J. Invertebr. Pathol. 132, 57–65.
7. Balsamo, P. J., Domingues, C. E. D. C., Silva-Zacarin, E. C. M. D., Gregorc, A., Irazusta, S. P., Salla, R. F., Costa, M. J., Abdalla, F. C., 2020. Impact of sublethal doses of thiamethoxam and *Nosema ceranae* inoculation on the hepato-nephrocitic system in young Africanized *Apis mellifera*. J. Apic. Res. 59, 350–361.
8. Bernklau, E., Bjostad, L., Hogeboom, A., Carlisle, A., Arathi, H. S., 2019. Dietary Phytochemicals, Honey Bee Longevity and Pathogen Tolerance. Insects 10, 14.
9. Borsuk, G., Paleolog, J., Olszewski, K., Strachecka, A., 2013. Laboratory assessment of the effect of nanosilver on longevity, sugar syrup ingestion, and infection of honeybees with *Nosema* spp. Med. Weter. 69, 730–732.
10. Braglia, C., Alberoni, D., Porrini, M. P., Garrido, P. M., Baffoni, L., Di Gioia, D., 2021. Screening of Dietary Ingredients against the Honey Bee Parasite *Nosema ceranae*. Pathogens 10, 1117.
11. Bravo, J., Carbonell, V., Sepúlveda, B., Delporte, C., Valdovinos, C. E., Martín-Hernández, R., Higes, M., 2017. Antifungal activity of the essential oil obtained from *Cryptocarya alba* against infection in honey bees by *Nosema ceranae*. J. Invertebr. Pathol. 149, 141–147.
12. Buczek, K., Deryło, K., Kutyła, M., Rybicka-Jasińska, K., Gryko, D., Borsuk, G., Rodzik, B., Trytek, M., 2020. Impact of Protoporphyrin Lysine Derivatives on the Ability of *Nosema ceranae* Spores to Infect Honeybees. Insects 11, 504.
13. Castelli, L., Balbuena, S., Branchiccela, B., Zunino, P., Liberti, J., Engel, P., & Antúnez, K., 2021. Impact of Chronic Exposure to Sublethal Doses of Glyphosate on Honey Bee Immunity, Gut Microbiota and Infection by Pathogens. Microorganisms 9, 845.

14. Cho, R. M., Kogan, H. V., Elikan, A. B., Snow, J. W., 2022. Paromomycin Reduces *Vairimorpha* (*Nosema*) *ceranae* Infection in Honey Bees but Perturbs Microbiome Levels and Midgut Cell Function. Microorganisms 10, 1107.
15. Clinch, P. G., Faulke, J., 1977. Effect of population density and storage temperature on the longevity of caged worker honey bees. New Zealand J. Exp. Agric. Res. 5, 67–69.
16. Czekońska, K., 2007. Influence of carbon dioxide on *Nosema apis* infection of honeybees (*Apis mellifera*). J. Invertebr. Pathol. 95, 84–86.
17. Diaz, T., del-Val, E., Ayala, R., Larsen, J., 2019. Alterations in honey bee gut microorganisms caused by *Nosema* spp. and pest control methods. Pest Manag. Sci. 75, 835–843.
18. Dosselli, R., Grassl, J., Carson, A., Simmons, L. W., Baer, B., 2016. Flight behaviour of honey bee (*Apis mellifera*) workers is altered by initial infections of the fungal parasite *Nosema apis*. Sci. Rep. 6, 36649.
19. Doublet, V., Natsopoulou, M. E., Zschiesche, L., Paxton, R. J., 2015. Within-host competition among the honey bees pathogens *Nosema ceranae* and Deformed wing virus is asymmetric and to the disadvantage of the virus. J. Invertebr. Pathol. 124, 31–34.
20. Dussaubat, C., Brunet, J.-L., Higes, M., Colbourne, J. K., Lopez, J., Choi, J.-H., Martín-Hernández, R., Botías, C., Cousin, M., McDonnell, C., Bonnet, M., Belzunces, L. P., Moritz, R. F. A., Le Conte, Y., Alaux, C., 2012. Gut Pathology and Responses to the Microsporidium *Nosema ceranae* in the Honey Bee *Apis mellifera*. PLoS ONE 7, e37017.
21. Eiri, D. M., Suwannapong, G., Endler, M., Nieh, J. C., 2015. *Nosema ceranae* Can Infect Honey Bee Larvae and Reduces Subsequent Adult Longevity. PLoS ONE 10, e0126330.
22. El Khoury, S., Rousseau, A., Lecoeur, A., Cheaib, B., Bouslama, S., Mercier, P.-L., Demey, V., Castex, M., Giovenazzo, P., Derome, N., 2018. Deleterious Interaction Between Honeybees (*Apis mellifera*) and its Microsporidian Intracellular Parasite *Nosema ceranae* Was Mitigated by Administrating Either Endogenous or Allochthonous Gut Microbiota Strains. Front. Ecol. Evol. 6, 58.
23. Faita, M. R., Cardozo, M. M., Amandio, D. T. T., Orth, A. I., Nodari, R. O., 2020. Glyphosate-based herbicides and *Nosema* sp. microsporidia reduce honey bee (*Apis mellifera* L.) survivability under laboratory conditions. J. Apic. Res. 59, 332–342.
24. Ferguson, J. A., Northfield, T. D., Lach, L., 2018. Honey Bee (*Apis mellifera*) Pollen Foraging Reflects Benefits Dependent on Individual Infection Status. Microbial Ecol. 76, 482–491.
25. Fontbonne, R., Garnery, L., Vidau, C., Aufauvre, J., Texier, C., Tchamitchian, S., Alaoui, H. E., Brunet, J.-L., Delbac, F., Biron, D. G., 2013. Comparative susceptibility of three Western honeybee taxa to the microsporidian parasite *Nosema ceranae*. Infect. Genet. Evol. 17, 188–194.
26. Forsgren, E., Fries, I., 2010. Comparative virulence of *Nosema ceranae* and *Nosema apis* in individual European honey bees. Vet. Parasitol. 170, 212–217.
27. Gajda, A. M., Mazur, E. D., Bober, A. M., Czopowicz, M., 2021. *Nosema ceranae* Interactions with *Nosema apis* and Black Queen Cell Virus. Agriculture 11, 963.
28. Giacomini, J. J., Leslie, J., Tarpy, D. R., Palmer-Young, E. C., Irwin, R. E., Adler, L. S., 2018. Medicinal value of sunflower pollen against bee pathogens. Sci. Rep. 8, 14394.

29. Glavinic, U., Blagojevic, J., Ristanic, M., Stevanovic, J., Lakic, N., Mirilovic, M., Stanimirovic, Z., 2022. Use of Thymol in *Nosema ceranae* Control and Health Improvement of Infected Honey Bees. Insects 13, 574.
30. Glavinic, U., Rajkovic, M., Vunduk, J., Vejnovic, B., Stevanovic, J., Milenkovic, I., Stanimirovic, Z., 2021. Effects of *Agaricus bisporus* Mushroom Extract on Honey Bees Infected with *Nosema ceranae*. Insects 12, 915.
31. Glavinic, U., Stankovic, B., Draskovic, V., Stevanovic, J., Petrovic, T., Lakic, N., Stanimirovic, Z., 2017. Dietary amino acid and vitamin complex protects honey bee from immunosuppression caused by *Nosema ceranae*. PLoS ONE 12, e0187726.
32. Glavinic, U., Stevanovic, J., Ristanic, M., Rajkovic, M., Davitkov, D., Lakic, N., Stanimirovic, Z., 2021. Potential of Fumagillin and *Agaricus blazei* Mushroom Extract to Reduce *Nosema ceranae* in Honey Bees. Insects 12, 282.
33. Grassl, J., Holt, S., Cremen, N., Peso, M., Hahne, D., Baer, B., 2018. Synergistic effects of pathogen and pesticide exposure on honey bee (*Apis mellifera*) survival and immunity. J. Invertebr. Pathol. 159, 78–86.
34. Gregorc, A., Jurišić, S., Sampson, B., 2019. Hydroxymethylfurfural Affects Caged Honey Bees (*Apis mellifera carnica*). Diversity 12, 18.
35. Gregorc, A., Silva-Zacarin, E. C. M., Carvalho, S. M., Kramberger, D., Teixeira, E. W., Malaspina, O., 2016. Effects of *Nosema ceranae* and thiametoxam in *Apis mellifera*: A comparative study in Africanized and Carniolan honey bees. Chemosphere 147, 328–336.
36. Hendriksma, H. P., Bain, J. A., Nguyen, N., Nieh, J. C., 2020. Nicotine does not reduce *Nosema ceranae* infection in honey bees. Insect. Soc. 67, 249–259.
37. Higes, M., García-Palencia, P., Martín-Hernández, R., Meana, A., 2007. Experimental infection of *Apis mellifera* honeybees with *Nosema ceranae* (Microsporidia). J. Invertebr. Pathol. 94, 211–217.
38. Higes, M., Martín-Hernández, R., García-Palencia, P., Marín, P., Meana, A., 2009. Horizontal transmission of *Nosema ceranae* (Microsporidia) from worker honeybees to queens (*Apis mellifera*). Environ. Microbiol. Rep. 1, 495–498.
39. Kang, E. J., Choi, Y. S., Byoun, G.-H., Lee, M.-Y., Kim, H. K., Lee, M. L., 2016. The Effect of CH4 on Life Span and *Nosema* Infection Rate in Honeybees, *Apis mellifera* L. J. Apic. 31, 331.
40. Lecocq, A., Jensen, A. B., Kryger, P., Nieh, J. C., 2016. Parasite infection accelerates age polyethism in young honey bees. Sci. Rep. 6, 22042.
41. Lee, M.-L., Byoun, G.-H., Lee, M.-Y., Choi, Y.-S., Kim, H.-K., 2015. The Effect of Temperature, Yellow Sand, and Acid Rain on Life Span and *Nosema* Infection Rate in Honeybees, *Apis mellifera* L. J. Apic. 30, 269.
42. Li, W., Evans, J. D., Huang, Q., Rodríguez-García, C., Liu, J., Hamilton, M., Grozinger, C. M., Webster, T. C., Su, S., Chen, Y. P., 2016. Silencing the Honey Bee (*Apis mellifera*) Naked Cuticle Gene (nkd) Improves Host Immune Function and Reduces *Nosema ceranae* Infections. Appl. Environ. Microbiol. 82, 6779–6787.
43. MacInnis, C. I., Keddie, B. A., Pernal, S. F., 2020. *Nosema ceranae* (Microspora: Nosematidae): A Sweet Surprise? Investigating the Viability and Infectivity of *N. ceranae* Spores Maintained in Honey and on Beeswax. J. Econ. Entomol. 113, 2069–2078.

44. MacInnis, C. I., Keddie, B. A., Pernal, S. F., 2021. Honey bees with a drinking problem: Potential routes of *Nosema ceranae* spore transmission. Parasitology 149, 573–580.
45. Maistrello, L., Lodesani, M., Costa, C., Leonardi, F., Marani, G., Caldon, M., Mutinelli, F., Granato, A., 2008. Screening of natural compounds for the control of *Nosema* disease in honeybees (*Apis mellifera*). Apidologie 39, 436–445.
46. Malone, L. A., Gatehouse, H. S., 1998. Effects of *Nosema apis* Infection on Honey Bee (*Apis mellifera*) Digestive Proteolytic Enzyme Activity. J. Invertebr. Pathol. 71, 169–174.
47. Malone, L. A., Giacon, H. A., Newton, M. R., 1995. Comparison of the responses of some New Zealand and Australian honey bees (*Apis mellifera* L) to *Nosema apis* Z. Apidologie 26, 495–502.
48. Martín-Hernández, R., Botías, C., Barrios, L., Martínez-Salvador, A., Meana, A., Mayack, C., Higes, M., 2011. Comparison of the energetic stress associated with experimental *Nosema ceranae* and *Nosema apis* infection of honeybees (*Apis mellifera*). Parasitol. Res. 109, 605–612.
49. Mayack, C., Naug, D., 2009. Energetic stress in the honeybee *Apis mellifera* from *Nosema ceranae* infection. J. Invertebr. Pathol. 100, 185–188.
50. Moffet, J.O., Lawson, F.A., 1975. Effect of *Nosema*-Infection on $O_2$ Consumption by Honey Bees. J. Econ. Entomol. 68, 627–629.
51. Nanetti, A., Ugolini, L., Cilia, G., Pagnotta, E., Malaguti, L., Cardaio, I., Matteo, R., Lazzeri, L., 2021. Seed Meals from *Brassica nigra* and *Eruca sativa* Control Artificial *Nosema ceranae* Infections in *Apis mellifera*. Microorganisms 9, 949.
52. Naree, S., Ellis, J. D., Benbow, M. E., Suwannapong, G., 2021. The use of propolis for preventing and treating *Nosema ceranae* infection in western honey bee (*Apis mellifera* Linnaeus, 1787) workers. J. Apic. Res. 60, 686–696.
53. Pajuelo, A. G., Torres, C., Bermejo, F. J. O., 2008. Colony losses: A double blind trial on the influence of supplementary protein nutrition and preventative treatment with fumagillin against *Nosema ceranae*. J. Apic. Res. 47, 84–86.
54. Paxton, R. J., Klee, J., Korpela, S., Fries, I., 2007. *Nosema ceranae* has infected *Apis mellifera* in Europe since at least 1998 and may be more virulent than *Nosema apis*. Apidologie 38, 558–565.
55. Peghaire, E., Moné, A., Delbac, F., Debroas, D., Chaucheyras-Durand, F., El Alaoui, H., 2020. A *Pediococcus* strain to rescue honeybees by decreasing *Nosema ceranae*- and pesticide-induced adverse effects. Pestic. Biochem. Phys. 163, 138–146.
56. Pettis, J. S., Lichtenberg, E. M., Andree, M., Stitzinger, J., Rose, R., vanEngelsdorp, D., 2013. Crop Pollination Exposes Honey Bees to Pesticides Which Alters Their Susceptibility to the Gut Pathogen *Nosema ceranae*. PLoS ONE 8, e70182.
57. Porrini, L. P., Porrini, M. P., Garrido, M. P., Müller, F., Arrascaeta, L., Fernández Iriarte, P. J., Eguaras, M. J., 2020. Infectivity and virulence of *Nosema ceranae* (Microsporidia) isolates obtained from various *Apis mellifera* morphotypes. Entomol. Exp. Appl. 168, 286–294.
58. Ptaszyńska, A. A., Borsuk, G., Mułenko, W., Olszewski, K., 2013. Impact of ethanol on *Nosema* spp. infected bees. Med. Weter. 69, 736-740.
59. Ptaszyńska, A. A., Borsuk, G., Mułenko, W., Wilk, J., 2016. Impact of vertebrate probiotics on honeybee yeast microbiota and on the course of nosemosis. Med. Weter. 72, 430–434.

60. Ptaszyńska, A. A., Paleolog, J., Borsuk, G., 2016. *Nosema ceranae* Infection Promotes Proliferation of Yeasts in Honey Bee Intestines. PLoS ONE 11, e0164477.
61. Rinderer, T.E., Elliott, K.E., 1977. Worker Honey Bee Response to Infection with *Nosema apis*: Influence of Diet. J. Econ. Entomol. 70, 431-433.
62. Rinderer, T. E., Sylvester, H. A., 1978. Variation in Response to *Nosema apis*, Longevity, and Hoarding Behavior in a Free-Mating Population of the Honey Bee. Ann. Entomol. 71, 372–374.
63. Roussel, M., Villay, A., Delbac, F., Michaud, P., Laroche, C., Roriz, D., El Alaoui, H., Diogon, M., 2015. Antimicrosporidian activity of sulphated polysaccharides from algae and their potential to control honeybee nosemosis. Carbohydr. Polym. 133, 213–220.
64. Simone-Finstrom, M., Aronstein, K., Goblirsch, M., Rinkevich, F., De Guzman, L., 2018. Gamma irradiation inactivates honey bee fungal, microsporidian, and viral pathogens and parasites. J. Invertebr. Pathol. 153, 57–64.
65. Strachecka, A. J., Olszewski, K., Paleolog, J., 2015. Curcumin Stimulates Biochemical Mechanisms of *Apis mellifera* Resistance and Extends the Apian Life-Span. J. Apic. Sci. 59, 129–141.
66. Strachecka, A., Krauze, M., Olszewski, K., Borsuk, G., Paleolog, J., Merska, M., Chobotow, J., Bajda, M., Grzywnowicz, K., 2014. Unexpectedly strong effect of caffeine on the vitality of western honeybees (*Apis mellifera*). Biochem. (Mosc.) 79, 1192–1201.
67. Strachecka, A., Olszewski, K., Paleolog, J., 2016. Varroa treatment with bromfenvinphos markedly suppresses honeybee biochemical defence levels. Entomol. Exp. Appl. 160, 57–71.
68. Strachecka, A., Olszewski, K., Paleolog, J., Borsuk, G., Bajda, M., Krauze, M., Merska, M., Chobotow, J., 2014. Coenzyme Q10 treatments influence the lifespan and key biochemical resistance systems in the honeybee, *Apis mellifera*. Arch. Insect Biochem. Physiol. 86, 165–179.
69. Tesovnik, T., Zorc, M., Ristanić, M., Glavinić, U., Stevanović, J., Narat, M., Stanimirović, Z., 2020. Exposure of honey bee larvae to thiamethoxam and its interaction with *Nosema ceranae* infection in adult honey bees. Environ. Poll. 256, 113443.
70. Tlak Gajger, I., Vlainić, J., Šoštarić, P., Prešern, J., Bubnič, J., Smodiš Škerl, M. I., 2020. Effects on Some Therapeutical, Biochemical, and Immunological Parameters of Honey Bee (*Apis mellifera*) Exposed to Probiotic Treatments, in Field and Laboratory Conditions. Insects 11, 638.
71. Toplak, I., Jamnikar Ciglenečki, U., Aronstein, K., Gregorc, A, 2013. Chronic Bee Paralysis Virus and Nosema ceranae Experimental Co-Infection of Winter Honey Bee Workers (*Apis mellifera* L.). Viruses 5, 2282–2297.
72. Urbieta-Magro, A., Higes, M., Meana, A., Barrios, L., Martín-Hernández, R., 2019. Age and Method of Inoculation Influence the Infection of Worker Honey Bees (*Apis mellifera*) by *Nosema ceranae*. Insects 10, 417.
73. Valizadeh, P., Guzman-Novoa, E., Petukhova, T., Goodwin, P. H., 2021. Effect of feeding chitosan or peptidoglycan on *Nosema ceranae* infection and gene expression related to stress and the innate immune response of honey bees (*Apis mellifera*). J. Invertebr. Pathol. 185, 107671.

74. Van Den Heever, J. P., Thompson, T. S., Otto, S. J. G., Curtis, J. M., Ibrahim, A., Pernal, S. F., 2016. Evaluation of Fumagilin-B® and other potential alternative chemotherapies against *Nosema ceranae*-infected honeybees (*Apis mellifera*) in cage trial assays. Apidologie 47, 617–630.
75. Van Den Heever, J. P., Thompson, T. S., Otto, S. J. G., Curtis, J. M., Ibrahim, A., Pernal, S. F., 2016. The effect of dicyclohexylamine and fumagillin on *Nosema ceranae*-infected honey bee (*Apis mellifera*) mortality in cage trial assays. Apidologie 47, 663–670.
76. Van Der Vorst, E., Jacobs, F. J., 1980. Comparison of Colony-and Laboratory-Stored Pollen for Maintaining the Life of Caged Honeybees. J. Apic. Res. 19, 119–121.
77. Vidau, C., Panek, J., Texier, C., Biron, D. G., Belzunces, L. P., Le Gall, M., Broussard, C., Delbac, F., El Alaoui, H., 2014. Differential proteomic analysis of midguts from *Nosema ceranae*-infected honeybees reveals manipulation of key host functions. J. Invertebr. Pathol. 121, 89–96.
78. Woyciechowski, M., Moroń, D., 2009. Life expectancy and onset of foraging in the honeybee (*Apis mellifera*). Insect. Soc. 56, 193–201.
79. Youngsteadt, E., Appler, R. H., López-Uribe, M. M., Tarpy, D. R., Frank, S. D., 2015. Urbanization Increases Pathogen Pressure on Feral and Managed Honey Bees. PLoS ONE 10, e0142031.
