## Appendix B for "Nosemosis negatively affects honeybee survival: experimental and meta-analytic evidence"

[1]Doctoral School of Exact and Natural Sciences, Jagiellonian University, Kraków, Poland; [2]Institute of Environmental Sciences, Faculty of Biology, Jagiellonian University, Kraków, Poland; [3]Department of Integrative Biology, College of Biological Science, University of Guelph, Guelph, ON, Canada; [4]Biological Sciences Platform, Holland Bone and Joint Program, Sunnybrook Research Institute; Department of Laboratory Medicine & Pathobiology, University of Toronto, ON, Canada; [5]Institute of Systematics and Evolution of Animals of the Polish Academy of Sciences, Kraków, Poland; [6]Department of Plant Physiology, Breeding and Seed Science, Faculty of Agriculture and Economics, University of Agriculture in Krakow, Poland

Corresponding authors:

Monika Ostap-Chec,

Krzysztof Miler,

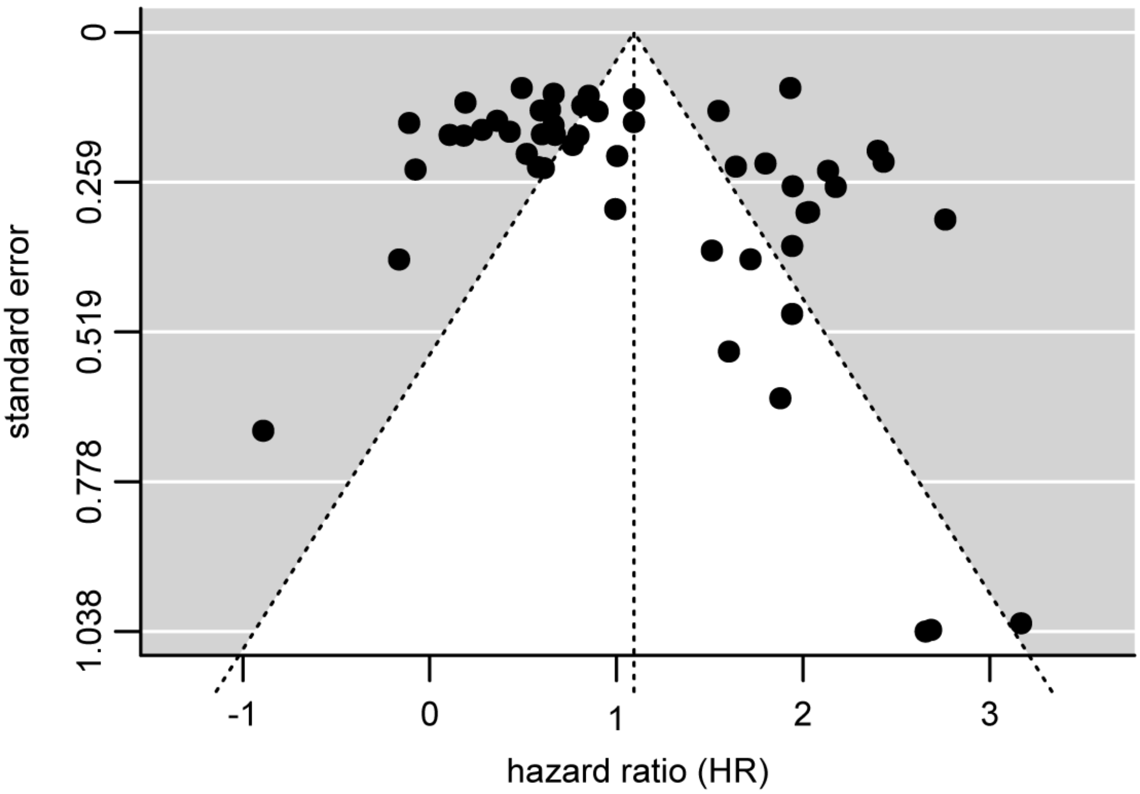


**Supplementary Figure 1**. Funnel plot (standard error plotted against hazard ratios (HRs)) for the studies included in the meta-analysis. Each dot represents one study.
